# Neutralizing antibodies targeting the SARS-CoV-2 receptor binding domain isolated from a naïve human antibody library

**DOI:** 10.1101/2021.01.07.425806

**Authors:** Benjamin N. Bell, Abigail E. Powell, Carlos Rodriguez, Jennifer R. Cochran, Peter S. Kim

**Affiliations:** Department of Molecular and Cellular Physiology, Stanford University School of Medicine, Stanford, CA 94305; Department of Biochemistry, Stanford University School of Medicine, Stanford, CA 94305; Stanford ChEM-H, Stanford University, Stanford, CA 94305; xCella Biosciences, Menlo Park, CA 94025; Department of Bioengineering, Stanford University, Stanford, CA 94305; Department of Chemical Engineering, Stanford University, Stanford, CA 94305; Chan Zuckerberg Biohub, San Francisco, CA 94158

**Keywords:** COVID-19, SARS-CoV-2, RBD, neutralizing antibody, yeast surface display

## Abstract

Infection with SARS-CoV-2 elicits robust antibody responses in some patients, with a majority of the response directed at the receptor binding domain (RBD) of the spike surface glycoprotein. Remarkably, many patient-derived antibodies that potently inhibit viral infection harbor few to no mutations from the germline, suggesting that naïve antibody libraries are a viable means for discovery of novel SARS-CoV-2 neutralizing antibodies. Here, we used a yeast surface-display library of human naïve antibodies to isolate and characterize three novel neutralizing antibodies that target the RBD: one that blocks interaction with angiotensin-converting enzyme 2 (ACE2), the human receptor for SARS-CoV-2, and two that target other epitopes on the RBD. These three antibodies neutralized SARS-CoV-2 spike-pseudotyped lentivirus with IC_50_ values as low as 60 ng/mL *in vitro*. Using a biolayer interferometry-based binding competition assay, we determined that these antibodies have distinct but overlapping epitopes with antibodies elicited during natural COVID-19 infection. Taken together, these analyses highlight how *in vitro* selection of naïve antibodies can mimic the humoral response *in vivo*, yielding neutralizing antibodies and various epitopes that can be effectively targeted on the SARS-CoV-2 RBD.

## Introduction

Over 84 million cases of Coronavirus Disease 2019 (COVID-19) and 1.8 million deaths worldwide in the past eleven months^1^ have been caused by the novel human pathogen SARS-CoV-2^2,3^. Since the beginning of the COVID-19 pandemic, extensive work has been devoted to characterizing the antibody response to SARS-CoV-2 in response to both infection^4–13^ and vaccination^14–20^. Analysis of antibody responses has revealed that patients elicit robust neutralizing antibodies targeting the SARS-CoV-2 spike^7,8,12,21,22^, a trimeric glycoprotein responsible for receptor recognition and host cell entry^2,23,24^. Specifically, the spike receptor binding domain (RBD), which binds the host cell receptor ACE2, has been shown to be an immunodominant region of the spike^6–9,12^. Despite being only 25 kDa in size, the RBD elicits antibodies that fall into several subclasses targeting distinct, but overlapping, epitopes^7,10^. Many of these subclasses contain neutralizing antibodies, demonstrating that multiple sites on the RBD are vulnerable to inhibition and raising the possibility that RBD-targeting antibodies neutralize viral entry via a variety of mechanisms of action.

Remarkably, many RBD antibodies described to date have high similarity to their germline antibody precursors, with low rates of somatic hypermutation^6–8,10^. This observation is encouraging for RBD- and spike-based vaccine approaches, as it suggests that neutralizing antibodies can readily be elicited in healthy individuals. This germline similarity also highlights the potential use of naïve antibody libraries as a promising source of novel therapeutic antibodies and for comparison of clones selected *in vitro* to those elicited by natural COVID-19 infection or vaccination.

In order to identify RBD-targeting antibodies with high similarity to the germline, here we employed an *in vitro* approach based on a yeast surface-display library of human naïve antibodies expressed in a single-chain variable fragment (scFv) format. We subsequently isolated and characterized three novel neutralizing antibodies: one (ABP18) that blocks binding of RBD to ACE2, and two (NBP10, NBP11) that do not. These three antibodies neutralize SARS-CoV-2 spike-pseudotyped lentivirus *in vitro* with IC_50_ (half-maximal inhibitory concentration) values of 0.06-13 μg/mL. When we performed binding competition experiments using biolayer interferometry (BLI), we found that ABP18, NBP10, and NBP11 have epitopes that are distinct but overlap with epitopes that are well-defined of antibodies elicited by natural COVID-19 infection. Taken together, these analyses demonstrate that *in vitro* selections of naïve antibodies can mimic the humoral response *in vivo* to identify neutralizing antibodies. *In vitro* selections therefore may constitute an important tool for both vaccine development and antibody discovery for SARS-CoV-2 therapeutics.

## Results

### A human naïve antibody library yields antibodies targeting the SARS-CoV-2 RBD

To isolate antibodies targeting the SARS-CoV-2 RBD, we panned a yeast surface-display library developed by xCella Biosciences, xEmplar™ (Materials and Methods), that contains ~1 × 10^9^ scFv sequences and has been used to identify antibodies against novel targets such as the immune checkpoint VISTA^25^. We performed a total of five rounds of selection that included both magnetic- and fluorescence-activated cell sorting to enrich for RBD-binding clones (Materials and Methods). In the final round, we also used biotinylated ACE2 labeled with Streptavidin-Alexa Fluor 647 to differentiate between scFvs that disrupted the ACE2/RBD interaction (ACE2-blocking population) and those that did not (non-blocking population; Fig. 1A). We identified a single unique sequence from the ACE2-blocking population (scFv ABP18) and three unique sequences from the non-blocking population (scFvs NBP10, NBP11, and NBP18); only ABP18, NBP10, and NBP11 were confirmed to bind RBD (SI Fig. 1A).

**Figure 1:**
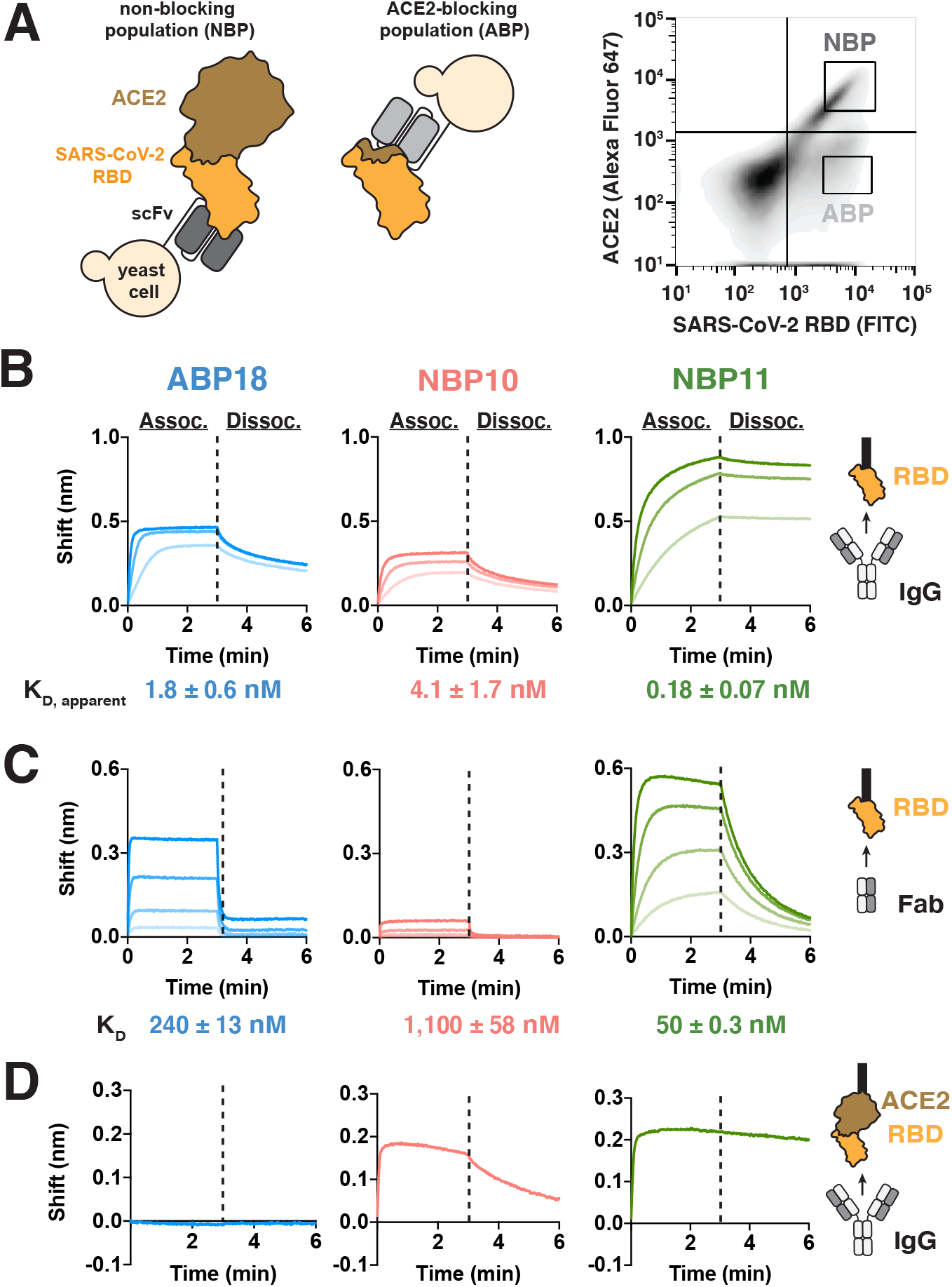
A human naïve antibody library yields antibodies targeting the SARS-CoV-2 RBD. **(A)** Left, selection scheme to isolate an ACE2-blocking population (ABP) and a non-blocking population (NBP) of antibodies. Right, flow cytometry panel of yeast display library with gates used to isolate the ABP and NBP. **(B-C)** Binding to SARS-CoV-2 RBD-coated BLI biosensors show substantial differences between **(B)** IgG antibodies (100, 33.3, and 11.1 nM) and **(C)** Fab antibody fragments (ABP18/NBP11: 500, 167, 56, 18 nM; NBP10: 720, 240, 80, 27 nM). Higher antibody concentrations are indicated with thicker lines. **(D)** Binding of 100 nM IgG antibodies to BLI biosensors coated with ACE2:RBD complex shows that NBP10 and NBP11 IgG retain binding while ABP18 shows no binding, consistent with ACE2-blocking activity.

We cloned the variable heavy chain and variable light chain sequences into human IgG1 plasmids (Materials and Methods) to further characterize ABP18, NBP10, and NBP11. We used BLI to confirm binding to RBD and to determine whether these antibodies compete with ACE2, as expected based on the sorting scheme. RBD-coated biosensors bound ABP18, NBP10, and NBP11 in IgG form (Fig. 1B), with low-nanomolar affinities calculated with a standard 1:1 binding model to simplify the bivalent interaction (Materials and Methods). To evaluate monovalent binding affinities, we processed the IgGs to Fab fragments using the endoprotease Lys-C (SI Fig. 1B). ABP18, NBP10, and NBP11 Fabs bound RBD (Fig. 1C) and exhibited substantially faster dissociation rates than the IgG forms (compare Fig. 1B,C), suggesting that the slow off-rates for IgG/RBD binding were influenced by the avidity of the antibody binding site. Consistent with the sequence and structural diversity among coronavirus RBDs^26^, enzyme-linked immunosorbent assays indicated that ABP18, NBP10, and NBP11 minimally bind RBDs from SARS-CoV-1 and MERS-CoV (SI Fig. 1C).

To confirm that our selection scheme differentiated between ACE2-blocking and non-blocking scFv clones, we employed BLI in two orientations to assess the ability of these antibodies to interact with RBD in the presence of ACE2 (SI Fig. 1D, E). When ACE2-coated biosensors were subsequently loaded with RBD, ABP18 showed no binding (Fig. 1D), while NBP10 and NBP11 IgG retained binding (Fig. 1D). In the reverse binding orientation using RBD-coated biosensors loaded with ACE2, ABP18 IgG binding was reduced when biosensors were preloaded with ACE2 (SI Fig. 1E); however, NBP10 and NBP11 IgG binding to RBD was unaffected by ACE2 (SI Fig. 1E), validating the ability of our selection scheme to distinguish between ACE2-blocking and non-blocking antibodies.

### RBD antibodies neutralize SARS-CoV-2 spike-pseudotyped lentivirus

Not all RBD-targeting antibodies neutralize SARS-CoV-2 infection^10,27–29^; further, antibodies isolated against the RBD alone may not effectively target the RBD in the context of the full-length spike trimer. We therefore tested the ability of ABP18, NBP10, and NBP11 IgG to neutralize SARS-CoV-2 pseudotyped lentivirus infection of ACE2-expressing HeLa cells^19,30^ (Fig. 2A). All three antibodies inhibited SARS-CoV-2 infection, with IC_50_ values ranging from 0.06 to 13 μg/mL (Fig. 2B,C).

**Figure 2:**
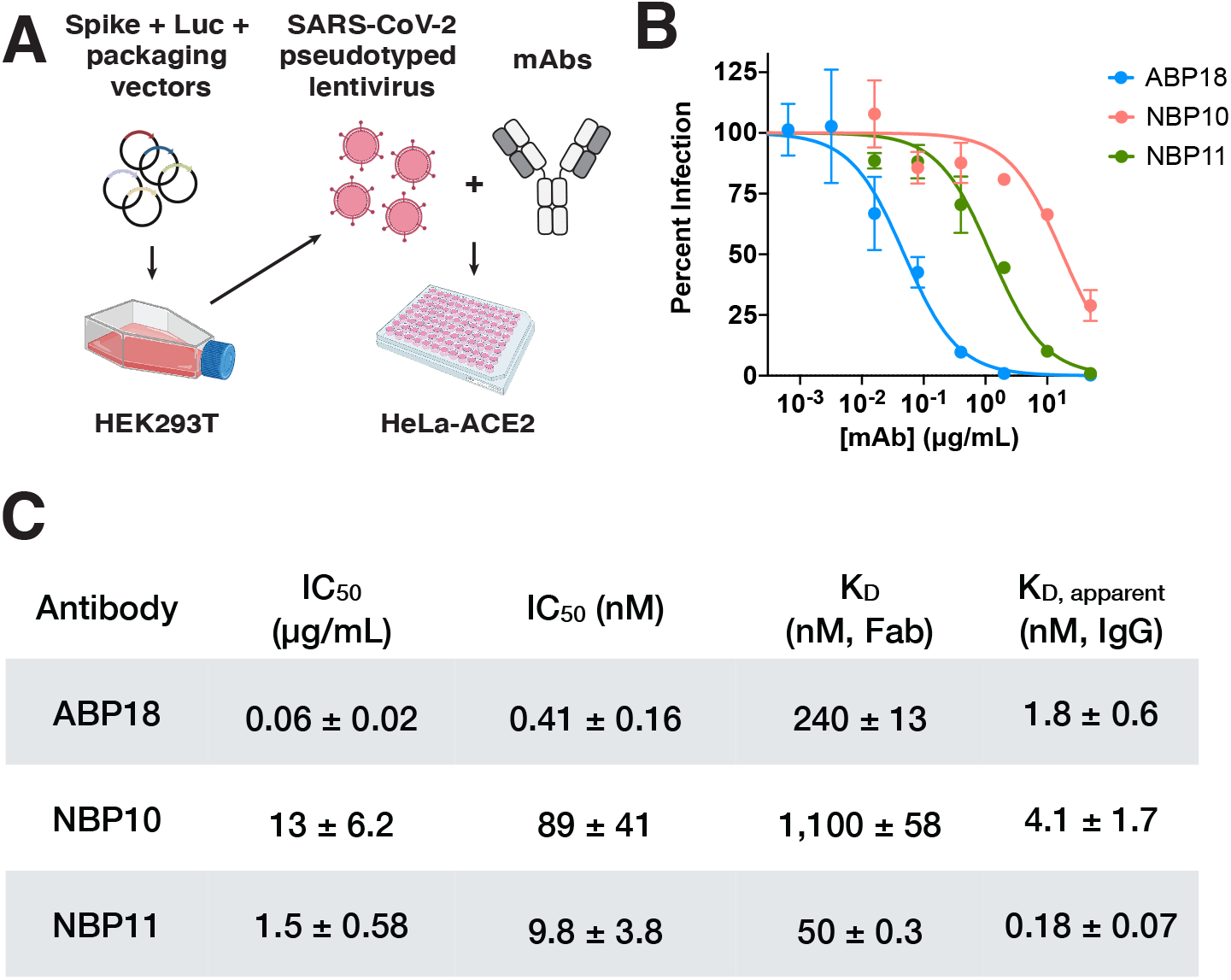
RBD antibodies neutralize SARS-CoV-2 pseudotyped lentivirus. **(A)** SARS-CoV-2 pseudotyped lentivirus produced in HEK293T cells encode firefly luciferase (Luc) and can infect HeLa cells stably expressing the ACE2 receptor (HeLa-ACE2)^19,20^. Half maximal inhibitory concentration (IC_50_) can be determined by titrating a monoclonal antibody (mAb) of interest. Figure created with BioRender.com. **(B-C)** ABP, NBP10, and NBP11 IgG **(B)** neutralize SARS-CoV-2 spike-pseudotyped lentivirus in vitro with **(C)** IC_50_ values (mean ± standard deviation) consistent with their true or apparent affinities. Representative inhibition curves are shown from repeated (n=3) experiments.

ABP18 IgG had an IC_50_ value of 0.41 nM (Fig. 2C), similar to its apparent affinity of 1.8 nM (Fig. 2C), while NBP10 and NBP11 IgG had IC_50_ values ~20- to 50-fold weaker than their apparent affinities (Fig. 2C). This difference may arise from (i) distinct mechanisms of viral neutralization employed by ABP18 and NBP10/NBP11 or (ii) differential epitope accessibility in the context of full-length spike or up/down conformations of the RBD^10,31^. Notably, all three antibodies have more potent IC_50_ values than their Fab binding affinities (Fig. 2C), suggesting that antibody avidity plays an important role in viral neutralization.

### Neutralizing RBD antibodies have distinct but overlapping epitopes with antibodies elicited by natural COVID-19 infection

Several classes of RBD antibodies have been described based on binding competition experiments and structure determination^6,10,13,19,27–29,32,33^. We therefore sought to map the epitopes of ABP18, NBP10, and NBP11 using a BLI-based binding competition assay. We expressed and purified (Materials and Methods) a panel of nine published RBD antibodies elicited by natural COVID-19 infection, four of which (CB6^33^, P2B-2F6^6^, EY6A^32^, CR3022^27^) have high-resolution co-crystal structures available. Five antibodies were included from Brouwer *et al*.^28^, four of which (COVA1-12, COVA2-04, COVA2-07, COVA2-15) have low-resolution negative-stain electron-micrograph reconstructions available^28^. We included the fifth antibody from Brouwer *et al*.^28^, COVA2-05, because it represents a unique epitope class (compared to COVA1-12, COVA2-04, COVA2-07, and COVA2-15) and shares the same antibody heavy chain germline as NBP10 and NBP11 (SI Table 1). However, no structure is available for COVA2-05. This nine-antibody panel represents a range of RBD epitopes and antibody sequences, generated *in vivo* via patient responses to infection, by which to compare the antibodies that we isolated *in vitro* here.

To evaluate how ABP18, NBP10, and NBP11 compete with antibodies in this panel for binding to RBD, we used BLI to establish a binding competition matrix (Materials and Methods). RBD-coated biosensors were bound to saturation by an antibody (the “loaded” antibody). These loaded-antibody/RBD-coated biosensors were then bound to antibodies in solution (the “associating” antibody). The binding of an associating antibody to biosensors coated with RBD and loaded with a competing antibody was compared to biosensors only coated with RBD and is reported as a fraction of maximal binding (Fig. 3A, SI Fig. 2A). Several ACE2-blocking antibodies (COVA1-12, COVA2-7, COVA2-4) displayed essentially the same competition binding profile (SI Fig. 2A), motivating us to use a reduced panel to analyze the binding profiles of ABP18, NBP10, and NBP11 (Fig. 3B).

**Figure 3:**
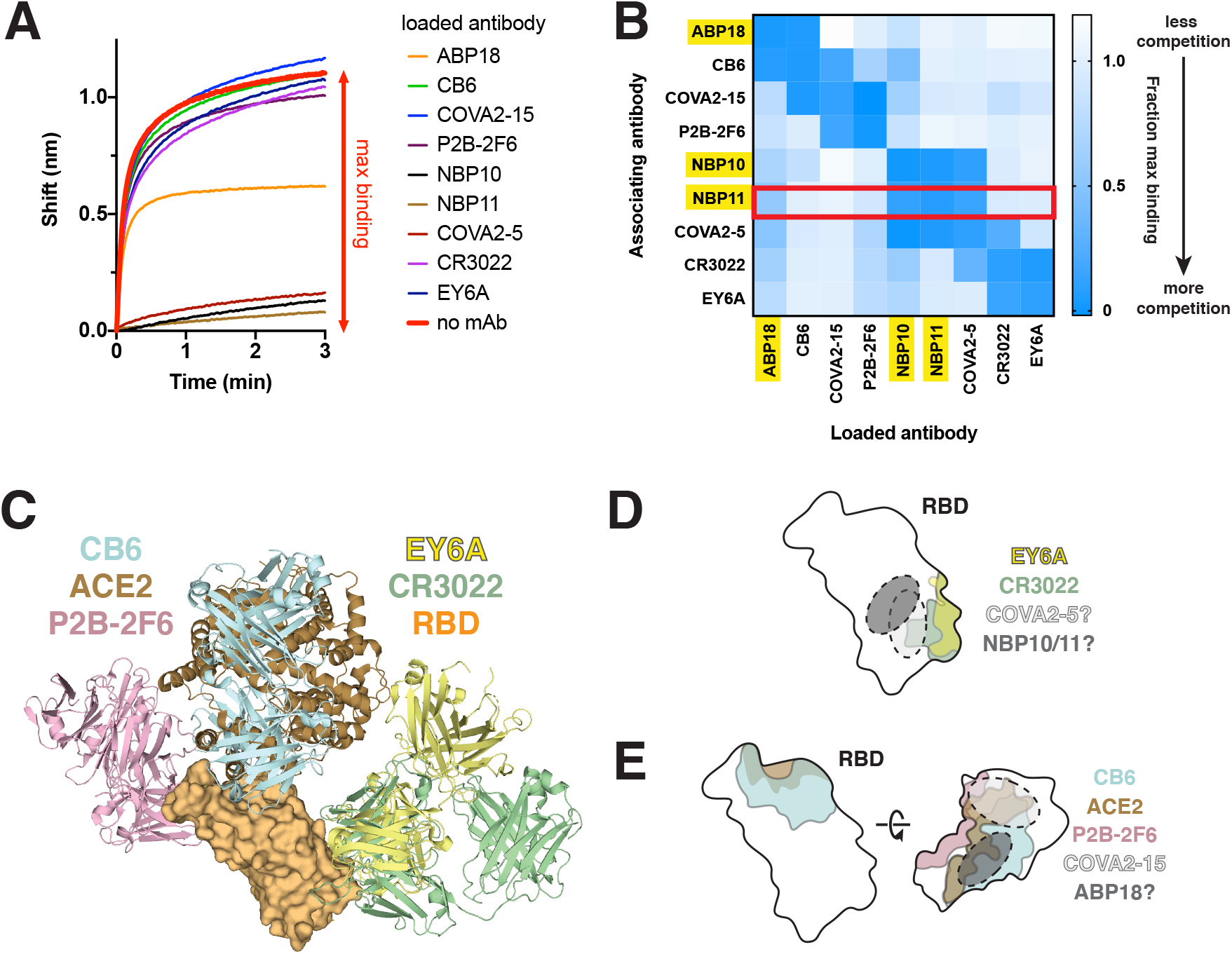
Neutralizing RBD antibodies have distinct but overlapping epitopes with antibodies elicited by natural COVID-19 infection. **(A)** Representative data from a BLI-based antibody competition assay with NBP11. RBD-coated biosensors were bound to saturation with an antibody (“loaded antibody”) before being dipped in a solution of NBP11 IgG (“associating antibody”). The shift observed when binding RBD-coated biosensors loaded with no antibody (red) is used to calculated fractional maximal binding. **(B)** Antibody competition matrix determined by BLI identifies approximate epitopes of ABP18, NBP10, and NBP11. Fractional maximal binding values for loaded antibodies (x-axis) dipped into solutions of associating antibodies (y-axis) are shown. Converted data for NBP11 in panel A are boxed in red. **(C)** Reported co-crystal structures of antibodies P2B-2F6 (PBD: 7BWJ), CB6 (7C01), CR3022 (6W41), EY6A (6ZER) against the RBD used for epitope mapping experiments; ACE2 (6M17) is shown for reference. **(D-E)** Approximate epitopes of **(D)** NBP10/11 and **(E)** ABP18 based on published X-ray crystal structures of indicated antibodies^6,27,32,33^. The approximate footprint of COVA2-15 is based on published negative-stain electron microscopy^28^.

NBP10 and NBP11 showed an identical pattern of binding competition (Fig. 3B), suggesting that they share similar epitopes, consistent with their common heavy chain germline, IGHV5-51 (SI Table 1). NBP10 and NBP11 competed with COVA2-05, but not CR3022 or EY6A (Fig. 3B). Interestingly, although EY6A and CR3022 have very similar epitopes^32^, COVA2-05 competed with CR3022, but not EY6A (Fig. 3B). We therefore conclude that NBP10, NBP11, COVA2-05, CR3022, and EY6A share overlapping but distinct epitopes. One notable difference between the epitopes of CR3022 and EY6A is that the complementarity determining region loop 1 of the CR3022 light chain extends beyond the EY6A light chain footprint (Fig. 3A), suggesting that this loop may be responsible for steric clashes between EY6A and NBP10/11. Combining our binding competition data and published structural data (Fig. 3C), we determined putative footprints for NBP10, NBP11, and COVA2-05 (Fig. 3D).

ABP18 demonstrated substantial competition with ACE2-blocking antibodies COVA1-12, COVA2-04, COVA2-07 (SI Fig. 2B), and CB6 (Fig. 3B), as expected. However, ABP18 did not compete with COVA2-15 or P2B-2F6 (Fig. 3B), which are other ACE2-competing antibodies. Based on the high-resolution X-ray crystal structure of P2B-2F6^6^ and the low-resolution negative-stain electron-micrograph reconstruction of COVA2-15^28^, we determined an approximate antibody footprint for ABP18 (Fig. 3E). Although the RBD makes up a small region of the spike trimer, these antibody competition results confirmed previous reports^7,10,29^ that the RBD contains several unique epitopes targeted by monoclonal antibodies. Further, the varied footprints and competition patterns even within the set of ACE2-competing antibodies tested here indicates that there are likely varied mechanisms of viral neutralization mediated by ACE2-blocking monoclonal antibodies.

## Discussion

Understanding the humoral responses to SARS-CoV-2 infection and vaccination is critical to efforts aimed at curtailing the COVID-19 pandemic and establishing effective correlates of protection. Accordingly, numerous patient-derived antibodies have been described, many of which target the RBD and have high similarity to germline antibody precursors. To exploit this similarity and to expand the repertoire of neutralizing antibodies, here we applied *in vitro* selections of a yeast surface-display library of human naïve antibodies to isolate and characterize three novel neutralizing antibodies that target the RBD: ABP18, NBP10, and NBP11 (Fig. 1). These antibodies neutralized SARS-CoV-2 spike-pseudotyped lentivirus *in vitro* (Fig. 2), targeting the ACE2-binding site (ABP18) and other RBD epitopes (NBP10 and NBP11) (Fig. 3). BLI-based epitope mapping experiments revealed that the epitopes of these novel antibodies are distinct but overlapping with antibodies elicited by natural COVID-19 infection (Fig. 3B). This work demonstrates that *in vitro* selections of naïve antibody repertoires can effectively mimic the antibody response *in vivo* in response to SARS-CoV-2 infection or vaccination, showcasing the value of this approach in rapidly isolating novel antibodies with therapeutic potential.

RBD-targeting antibodies bind numerous distinct, but overlapping, epitopes. We selected a panel of patient-derived RBD antibodies with well-defined epitopes and determined that none of them shared identical profiles with ABP18, NBP10, or NBP11 in binding competition experiments (Fig. 3B). Each of our three novel antibodies demonstrated strong competition with at least one antibody in the binding panel, which in turn had strong competition with additional antibodies (Fig. 3B), enabling us to delineate putative epitopes for ABP18, NBP10, and NBP11 (Fig. 3D,E). High-resolution structural information is needed to more precisely determine these antibody binding sites and their binding angles. Binding angles can be a critical factor in viral neutralization^34^; for example, antibodies such as CR3022 and EY6A complete strongly for binding the RBD, but only EY6A neutralizes SARS-CoV-2 entry^32^. Nevertheless, our results highlight how methods like BLI can quickly delineate classes of antibodies and provide low-resolution structural parameters for antibody:antigen interactions.

Antibodies with high similarity to the germline, and from diverse germline lineages, bind the RBD and can effectively neutralize SARS-CoV-2^7,10,29^. The antibodies described here also displayed high germline similarity, with low degrees of somatic hypermutation (SI Table 1). This observation is particularly promising for RBD- and spike-based vaccination strategies, as they may readily elicit neutralizing antibodies in patients presenting varied human antibody repertoires. There is well-documented enrichment of specific antibody germlines among antibodies elicited by SARS-CoV-2 infection, including IGHV1-24, IGVH1-69, IGHV3-30, IGHV3-53, and IGHV5-51^7,8,28,35^. The antibodies that we isolated here have sequences derived from these major germlines elicited in COVID-19 patients (SI Table 1). ABP18 is an IGHV1-69 antibody, a common germline regularly elicited by influenza infection^36^ and enriched among SARS-CoV-2 antibodies^28^. By contrast, NBP10 and NBP11 both utilize IGHV5-51, consistent with their shared epitope. NBP11 shares both heavy and light chain germlines (IGHV5-51 and IGKV1-5, respectively) with C110, a recently described neutralizing RBD antibody derived from a COVID-19 patient^29^. Interestingly, the epitope of P2B-2F6 overlaps with that of C110^10^, though P2B-2F6 does not compete with NBP11 (Fig. 3B), suggesting that C110 and NBP11 target distinct epitopes. This result further illustrates the diversity of near-germline antibodies that can effectively target the RBD, and underscores that even antibodies with the same germline precursors can target distinct RBD epitopes.

Taken together, these analyses highlight how *in vitro* selections of naïve antibodies can mimic the humoral response *in vivo* to identify neutralizing antibodies against novel human pathogens like SARS-CoV-2. Additionally, these novel RBD-targeting antibodies demonstrate the numerous distinct epitopes that can be effectively targeted on the SARS-CoV-2 RBD, bolstering COVID-19 therapeutic and vaccine efforts.

## Materials and Methods

### Yeast surface display selections

xEmplar™ is a proprietary ~1 × 10^9^ member scFv *Saccharomyces cerevisiae* yeast surface display library designed by xCella Biosciences. A total of five rounds of sorting were conducted to isolate scFv-displaying yeast that could bind the SARS-CoV-2 spike RBD and disrupt ACE2/RBD interactions. The antigens used for the selections were His-tagged RBD from Ray Biotech (Cat. # 230-30162), His-tagged and Avi-Tagged RBD from ACROBiosystems (Cat. # SPD-C82E9), and His-tagged and Avi-Tagged ACE2 from ACROBiosystems (Cat. # AC2-H82E6). Two initial rounds of magnetic-activated cell sorting were performed on the library using decreasing amounts of Ray Biotech RBD-coated His-Tag Dynabeads (Thermo Fisher). The resulting enriched library pool was subsequently sorted using a Sony SH800 Cell Sorter. Round 3 selected for variants that bound 200 nM RBD (Ray Biotech). Round 4 enriched for variants that bound 100 nM RBD (ACROBiosystems). In Round 5, the ACE2/RBD blocking sort, the enriched library was incubated with 100 nM RBD (Ray Biotech) for 1 h at room temperature, washed once using phosphate-buffered saline [pH 7.4] with 0.1% bovine serum albumin (PBSA), then incubated with secondary antibody for 15 min on ice to detect RBD binding. The resulting complex was washed twice with PBSA, incubated with 100 nM ACE2 (ACROBiosystems) for 30 min on ice, and washed once more with PBSA. Last, the cells were incubated with 1:1000 dilution of Streptavidin-Alexa Fluor 647 (Thermo Fisher Scientific Cat. # S32357) on ice for 15 min for detection of ACE2 binding.

Yeast scFv display levels were evaluated prior to sorting (data not shown) by incubating the yeast with a 1:1000 dilution of Chicken anti C-MYC Epitope Tag Primary Antibody from Exalpha Biologicals Inc. (Catalog No. ACMYC), followed by incubation with a 1:1000 dilution of Goat anti-Chicken IgY (H+L) Cross Adsorbed Secondary Antibody, Alexa Fluor Plus 647 from Thermo Fisher Scientific (Catalog No. A32933)

RBD antigen binding during selection rounds 3 and 5 was evaluated using a 1:1000 dilution of rabbit anti-His FITC secondary antibody (Bethyl Laboratories). RBD antigen binding during selection Round 4 was evaluated using a 1:1000 dilution of Streptavidin-Alexa Fluor 647 secondary (Jackson ImmunoResearch). Use of a different secondary reagent selected for true RBD binders and de-enriched the population for anti-His-tag secondary reagent binders (data not shown). In subsequent selections, such secondary reagent binders would give the false impression of being ACE2-blocking antibodies.

After Round 5, plasmid DNA was isolated from the sorted yeast populations using a Zymoprep Yeast Miniprep II Kit (Zymo Research). Stellar cells (Takara) were transformed with these polyclonal plasmid DNA mixtures and streaked to single colonies on LB agar plates supplemented with 100 μg/mL carbenicillin. Single-colony cultures were propagated in 2xYT medium supplemented with 100 μg/mL carbenicillin and the sequences of mini-prepped plasmids were analyzed to identify scFvs ABP18, NBP10, NBP11, and NBP18.

### Plasmid cloning and propagation

Variable heavy chain and light chain sequences were PCR amplified from yeast plasmid DNA and cloned into CMV/R IgG expression vectors using InFusion (Takara). Plasmid DNA for mammalian expression of the His-tagged SARS-CoV-2 RBD in a pCAGGS vector was kindly provided by Dr. Florian Krammer^37^. Plasmid DNA for expression of His-tagged SARS1 and MERS RBDs under control of a CMV promoter was a kind gift from Dr. John Pak. Plasmid DNA for expression of ACE2-huFc-Avi was previously described^20^. All plasmids were sequence confirmed via Sanger sequencing prior to mammalian transfection.

Plasmids were propagated overnight at 37 °C in Stellar cells (Takara) in 2xYT medium supplemented with 100 μg/mL carbenicillin (RBD and ACE2 plasmids) or 50 μg/mL kanamycin (IgG plasmids). Plasmids were isolated using a Machery Nagel maxi prep kit and filtered in a biosafety cabinet using a 0.22-μm filter prior to transfection.

### Recombinant protein expression and purification

IgGs, ACE2-huFc-Avi, and coronavirus RBDs were expressed and purified from Expi293F cells cultured in 33% Expi293 Expression / 66% FreeStyle Expression medium (Thermo Fisher Scientific) and grown in baffled polycarbonate shaking flasks (Triforest) at 37 °C and 8% CO_2_. Cells were grown to a density of ~3 x 10^6^/mL and transiently transfected using FectoPro transfection reagent (Polyplus). Briefly, 0.5 μg DNA per mL final transfection volume was added to culture medium (1/10 volume of final transfection) followed by FectoPro at a concentration of 1.3 μL per mL final transfection volume and incubated at room temperature for 10 min. Transfection mixtures were added to cells, which were then supplemented with D-glucose (4 g/L final concentration) and 2-propylpentanoic (valproic) acid (3 mM final concentration). Cells were harvested 3-5 days after transfection via centrifugation at 18,000 x *g* for 15 min. Cell culture supernatants were filtered using a 0.22-μm filter prior to purification.

Filtered Expi cell culture supernatants from ABP18, NBP10, and NBP11 IgG expressions were buffered with 1X PBS [pH 7.4] and purified using Protein A resin (Pierce). Supernatants were incubated with Protein A resin in batch and then resin/supernatant mixtures were added to glass chromatography columns. Resin was washed with ~10 column volumes of 1X PBS and IgGs were eluted with 100 mM glycine [pH 2.8] and immediately quenched with 1/10 volume 1 M Tris [pH 8.0]. ACE2 expressed with a C-terminal human IgG1 Fc and Avi tag was purified in the same way as IgGs.

Antibodies in the competition panel (see *Biolayer Interferometry*) were purified on a HiTrap MabSelect SuRe column (Cytiva) equilibrated in PBS [pH 7.4] on a GE ÄKTA Pure system. Filtered Expi cell culture supernatants were buffered with 1/10 volume 10x PBS and loaded at a flow rate of 3.5 mL/min. The column was subsequently equilibrated with 5 column volumes PBS [pH 7.4] before elution with 3 column volumes 100 mM glycine [pH 2.8] into 1/10^th^ volume of 1M Tris [pH 8.0]. The column was washed with 0.5 M NaOH with a minimum contact time of 15 min between purifications of different antibodies.

Coronavirus RBDs (SARS-CoV-2, SARS1, and MERS) were purified using HisPur Ni-NTA affinity resin (Thermo). Supernatants were diluted 1:1 with 10 mM imidazole / 1X PBS and incubated with Ni-NTA resin for at least 1 h at 4 °C while stirring. Resin/supernatant mixtures were then added to glass chromatography columns, washed with ~10 column volumes 10 mM imidazole / 1X PBS, and RBDs were eluted using 250 mM imidazole / 1X PBS. Elutions were concentrated using Amicon spin filters (MWCO 10 kDa; Millipore Sigma) and RBDs were subsequently loaded onto a GE Superdex S200 increase 10/300 GL column pre-equilibrated in 1X PBS on a GE ÄKTA Pure system. Protein-containing fractions were identified by A280 signal and/or SDS-PAGE, pooled, and stored at 4 °C or at −20 °C in 10% glycerol / 1X PBS until use.

### Isolation of Fab from IgG

Lys-C endopeptidase (Wako) was added at a final concentration of 4 μg/mL to purified IgG (2 mg/mL in PBS + 100 mM Tris-HCl [pH 8.0]). This mixture was incubated for 1-1.5 h at 37 °C with gentle shaking, then buffer exchanged to 50 mM NaOAc [pH 5.0] with Amicon spin filters (MWCO 10 kDa; Millipore Sigma). Fab was separated from undigested IgG and free Fc via cation exchange chromatography using a HiTrap SP HP column (Cytiva) over a 0-400 mM NaCl gradient in 50 mM NaOAc [pH 5.0]. Purified Fab was buffer exchanged into 1X PBS [pH 7.4] for use in BLI.

### ACE2-huFc-Avi biotinylation

Purified ACE2 expressed with a human IgG1 Fc domain and C-terminal Avi tag was site-specifically biotinylated using BirA-thioredoxin. BirA enzyme was expressed and purified as previously described^38^. Briefly, BirA was expressed recombinantly in *Escherichia coli* as a genetic fusion to thioredoxin and purified with HisPur Ni-NTA affinity resin (Thermo Fisher Scientific). BirA-thioredoxin was subsequently loaded onto a GE Superdex S200 increase 10/300 GL column pre-equilibrated in 1X PBS on a GE ÄKTA Pure system before being aliquoted and frozen with 10% glycerol.

To biotinylate purified ACE2, briefly, ACE2 was buffered with 50 mM bicine [pH 8.3], 50 mM biotin, and 10 mM ATP. BirA enzyme was added at a ratio of 1:200 w/w and incubated for 1 h at 37 °C before being buffer exchanged into PBS [pH 7.4] with a Zeba 7 kDa MWCO spin column (Thermo Fisher Scientific).^35^

### Enzyme-linked immunosorbent assay

Nunc 96-well Maxisorp plates (Thermo) were coated with 50 μL of a 2 μg/mL RBD solution from SARS-CoV-1, SARS-CoV-2, or MERS-CoV in 50 mM sodium bicarbonate [pH 8.75] at room temperature for 1 h. Plates were then washed 3 times with MilliQ ddH_2_O using an ELx 405 BioTek plate washer and blocked with 300 μL of Chonblock Blocking Buffer (Chondrex) at 4 °C overnight. Chonblock was manually removed from plates and plates were subsequently washed 3 times with PBST (1X PBS [pH 7.4] / 0.1% Tween-20). IgGs were diluted in PBST starting at 15 μg/mL with a 5-fold dilution series. Antibody dilutions were added to each well and incubated at room temperature for 1 h. Plates were washed 3 times with PBST and then incubated with a goat anti-human HRP (abcam ab7153) secondary, at a 1:10000 dilution in PBST for 1 h at room temperature. Following secondary incubation, plates were washed six times with PBST and developed by adding 50 μL of 1-Step^TM^ Turbo-TMB-ELISA Substrate Solution (Thermo Fisher Scientific 34022) per well. Plates were quenched by adding 50 μL of 2 M H2SO4 per well and absorbance at 450 nm was quantified and normalized for path length using a BioTek Synergy HT Microplate Reader.

### Biolayer interferometry (BLI)

His-tagged SARS-CoV-2 RBD (300 nM) was loaded on His1K biosensors (Pall ForteBio) to a load threshold of 0.4-0.6 nm. After a baseline step in assay buffer (PBS [pH 7.4], 0.1% bovine serum albumin, 0.05% Tween 20), ligand-loaded sensors were dipped into known concentrations of antibody for an association step and returned to the baseline well for dissociation. All samples in all experiments were baseline-subtracted to a well that loaded the tip with antibody, but did not go into sample, as a control for any buffer trends within the samples. Binding curves were fit using the “association then dissociation” equation in GraphPad Prism 8.4.1 to calculate KD.

For ACE2 binding competition, experiments were performed in one of two ways. (1) Biotinylated ACE2 (300 nM) was loaded on Streptavidin biosensors (Pall ForteBio) to a load threshold of 0.4-0.6 nm. After a baseline step in assay buffer, ACE2-coated biosensors were dipped in a 300 nM solution of RBD for 5 min before being dipped in a 100 nM solution of a given antibody. These results are reported in Fig. 1D. (2) RBD (300 nM) was loaded on His1K biosensors (Pall ForteBio) to a load threshold of 0.4-0.6 nm. After a baseline step in assay buffer, RBD-coated biosensors were dipped in a 300 nM solution of ACE2 or assay buffer for 5 min before being dipped in a 100 nM solution of a given antibody. These results are reported in Supplemental Fig. 1E.

For antibody binding competition assays, His-tagged SARS-CoV-2 RBD (300 nM) was loaded on His1K biosensors (Pall ForteBio) to a load threshold of 0.4-0.6 nm. After a baseline step in assay buffer, RBD-coated biosensors were dipped in a 500 nM solution of a particular “loaded” antibody for 4 min. “Loaded”-antibody/RBD-coated biosensors were then dipped in a 200 nM solution of a particular “associating” antibody for 3 min. The detected shift in nanometers of the “associating” antibody at 3 min in the absence of any “loaded” antibody was used as the maximal binding value. Values for the “loaded” antibody are reported as a fraction of this value. All BLI data were baseline-subtracted based on RBD-coated biosensors dipped into buffer for the “loading” and “associating” steps.

### Production of SARS-COV-2 pseudotyped lentivirus and viral neutralization assays

SARS-CoV-2 spike-pseudotyped lentivirus was produced as described^20^. Briefly, 6 x 10^6^ HEK293T cells cultured in D10 medium (DMEM, 10% fetal bovine serum (Gemini Bio-Products), 2 mM L-glutamine, 1% penicillin/streptomycin, and 10 mM HEPES [pH 7.0]) were transfected at ~50-70% confluency using calcium phosphate transfection^20^ with 5 lentiviral production plasmids: pHAGE_Luc packaging vector (10 μg), full-length SARS-CoV-2 spike (3.4 μg), HDM-Hgpm2 (2.2 μg), PRC-CMV-Rev1b (2.2 μg), and HDM-tat1b (2.2 μg)^30^. Filter-sterilized plasmids were added to 0.5 mL sterile water followed by addition of 0.5 mL 2X HEPES-buffered saline [pH 7.0] (Alfa Aesar) and then dropwise addition of 100 μL 2.5 M CaCl2. Transfections were incubated at room temperature for 20 min and then added dropwise to plated cells. Medium was exchanged ~18-24 h post transfection. Virus was harvested ~72 h post transfection via centrifugation at 300 x *g* for 5 min followed by filtration through a 0.45-μm filter. Viruses were aliquoted and stored at −80 °C.

Neutralization assays were performed as described^19,20^. HeLa cells overexpressing ACE2^19^ were plated at 5,000 cells per well in clear-bottom white-walled 96-well plates 1 day prior to infection in D10 medium. Purified monoclonal antibodies were filtered using a 0.22-μm filter and then diluted using D10 medium. Virus was diluted in D10 medium and supplemented with polybrene (5 μg/mL final), added to inhibitor dilutions, and inhibitor/virus were incubated at 37 °C for 1 h. Medium was removed from HeLa/ACE2 cells and replaced with virus/inhibitor mixtures which were then incubated for 48 h at 37 °C. Cells were lysed via addition of BriteLite (Perkin Elmer) and luminescence signal from wells was determined using a BioTek plate reader.

Data from neutralization assays were processed using GraphPad Prism 8.4.1. Relative luminescence units from virus-only wells and cells-only wells were averaged, and RLU values from inhibitor neutralization curves were normalized to these averages (100% infection for virus only and 0% infection for cells only). Normalized infectivity values were then plotted as a function of inhibitor concentration and fit with a non-linear 3-parameter inhibitor curve to obtain IC_50_ values.

## Key abbreviations

ACE2: angiotensin-converting enzyme 2
BLI: biolayer interferometry
COVID-19: Coronavirus Disease 2019
RBD: receptor-binding domain
scFv: single-chain variable fragment

## Acknowledgments

We thank the Kim Lab for critical reading of this manuscript. The CMV/R expression vectors were received from the NIH AIDS Reagent Program. We thank Dr. Jesse Bloom, Kate Crawford, Dr. Dennis Burton, and Dr. Deli Huang for sharing plasmids, cells, and invaluable advice for implementation of the spike-pseudotyped lentiviral neutralization assay. This work was supported by the National Science Foundation Graduate Research Fellowship Program (to BNB), the Stanford Maternal and Child Health Research Institute postdoctoral fellowship (to AEP), the Virginia and D. K. Ludwig Fund for Cancer Research (to PSK), the Chan Zuckerberg Biohub (to PSK), and the Frank Quattrone and Denise Foderaro Family Research Fund (to PSK).

## Author Contributions

**Benjamin N. Bell**: Investigation (lead); Resources; Writing-original draft; Writing-review and editing; Visualization

**Abigail E. Powell**: Investigation; Resources; Writing-review and editing

**Carlos Rodriguez**: Investigation; Resources; Writing-review and editing

**Jennifer R. Cochran**: Supervision; Writing-review and editing

**Peter S. Kim**: Supervision; Writing-review and editing

---

**Supplementary Figure 1:**
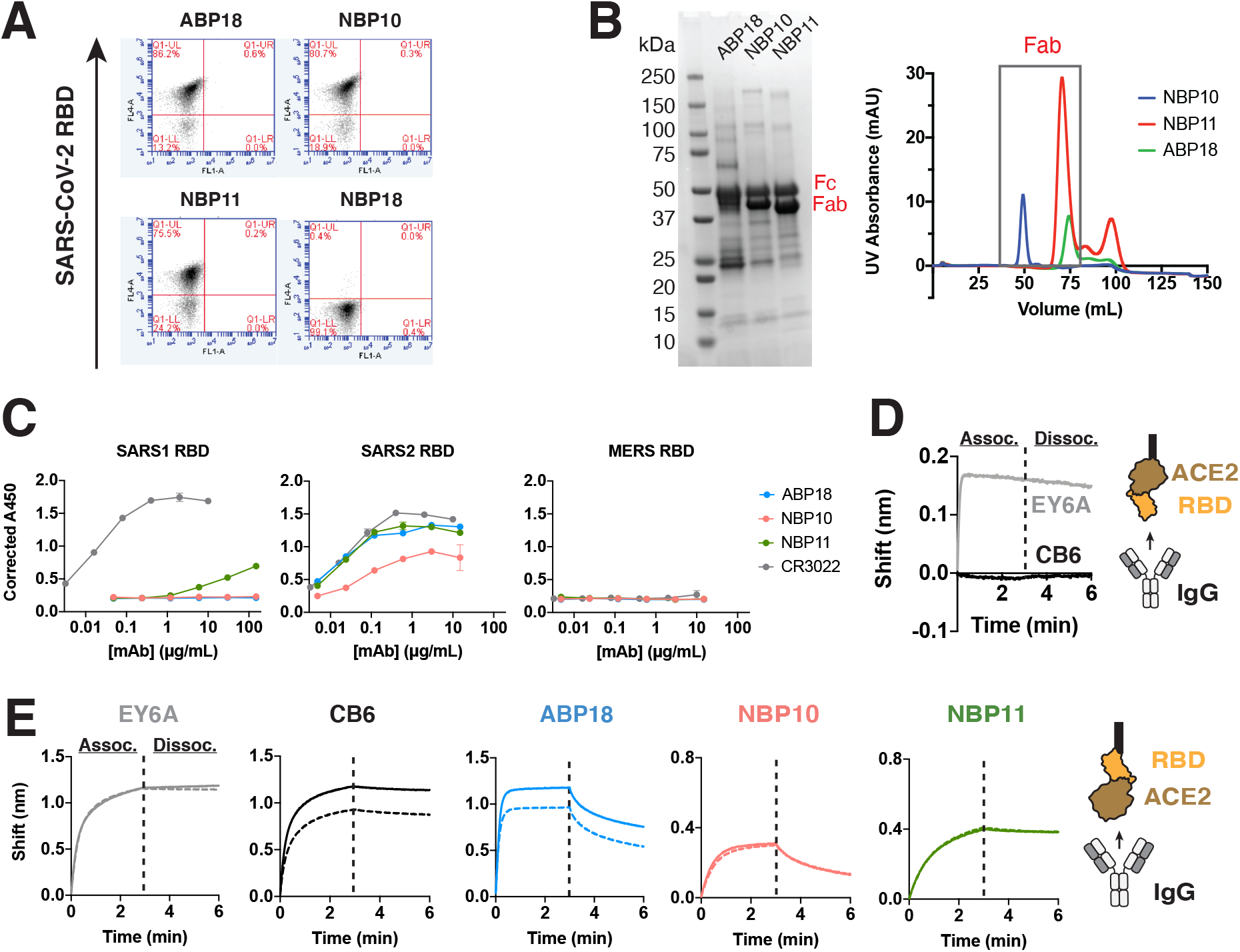
Extended data related to Figure 1. **(A)** Flow cytometry of four unique sequences isolated from selections of the xEmplar™ yeast surface-display library. Clones ABP18, NBP10, and NBP11 bind the SARS-CoV-2 RBD, but NBP18 does not. **(B)** ABP18, NBP10, and NBP11 Fabs were isolated via Lys-C cleavage of purified IgG (see Materials and Methods). Left, Coomassie staining of digested IgG samples. Right, corresponding ion exchange traces used to isolate Fabs. **(C)** Isolated RBD antibodies have minimal cross-reactivity to other human pandemic coronaviruses, as revealed by ELISA of CR3022 IgG and newly identified IgG antibodies to RBDs from human coronaviruses SARS-CoV-1, SARS-CoV-2, and MERS-CoV. Mean ± standard deviation is shown. **(D)** Binding of 100 nM IgG control antibodies to BLI biosensors coated with ACE2:RBD complex shows that EY6A IgG retains binding while CB6 shows no binding, consistent with ACE2-blocking activity. **(E)** BLI indicates that binding of 100 nM IgG antibodies to RBD-coated biosensors (solid line) is reduced when human ACE2 is preloaded on the biosensors (dashed line) only in the case of ABP18, consistent with ACE2-blocking activity.

**Supplementary Figure 2:**
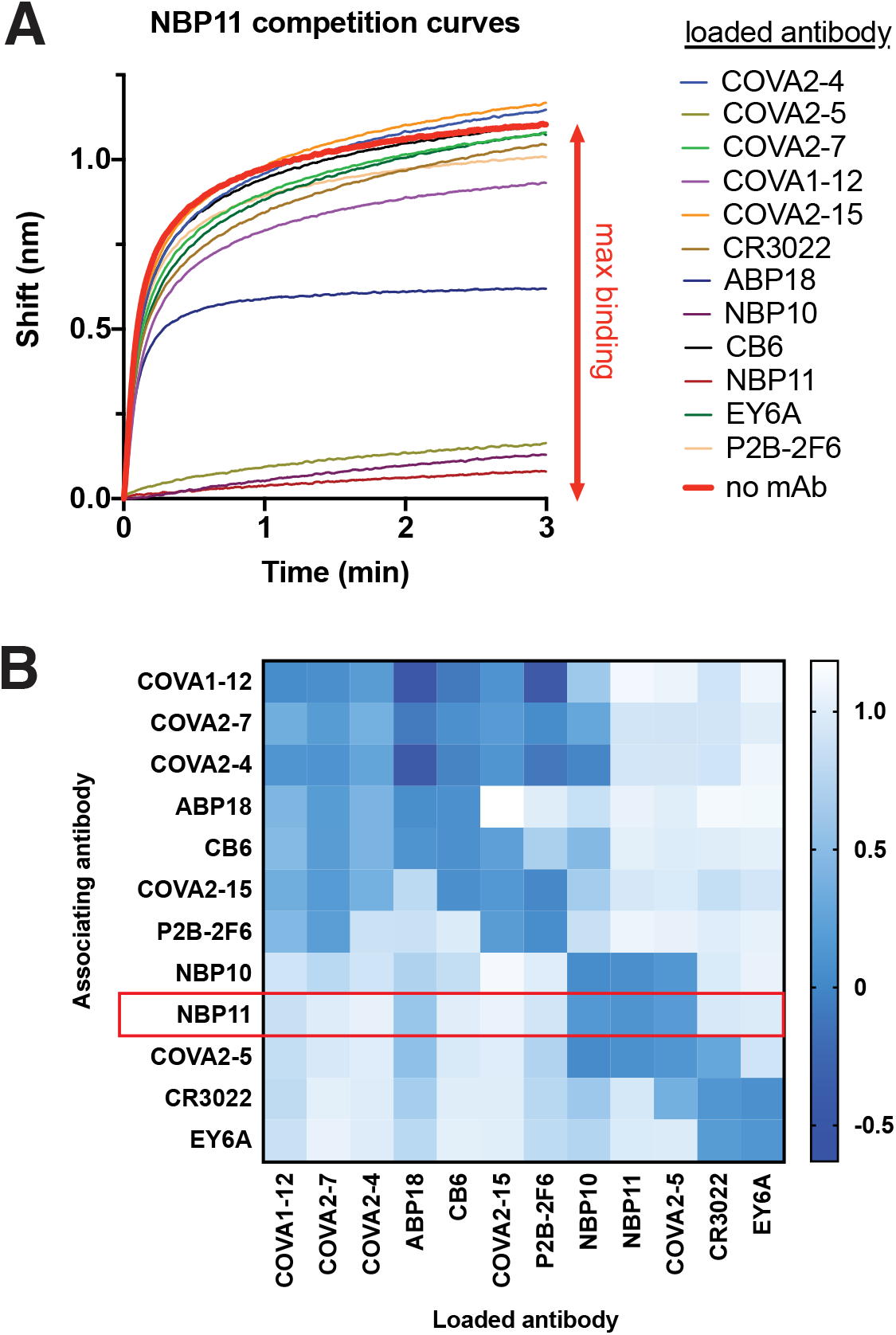
Extended data related to Figure 3. **(A)** BLI binding curves of 100 nM NBP11 IgG to RBD-coated biosensors loaded with an extended antibody panel. The shift observed when binding RBD-coated biosensors loaded with no antibody (red) is used to calculated fractional maximal binding. **(B)** Complete antibody competition matrix determined via BLI. RBD-coated biosensors were loaded to saturation with an antibody; subsequent binding to an associating antibody is reported as a fraction of binding to RBD-coated biosensors without loaded antibody. Converted data from panel A are boxed in red.

**Supplementary Table 1:**
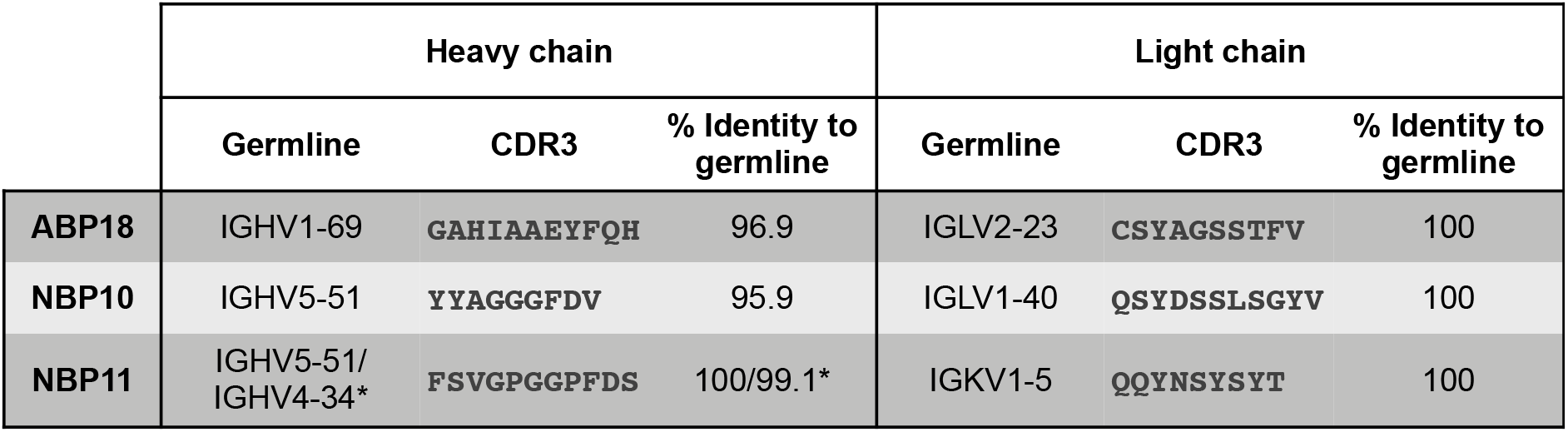
Sequence characteristics of ABP18, NBP10, and NBP11 antibodies. Isolated RBD antibodies represent germlines elicited by natural COVID-19 infection and have high sequence similarity to the germline. *The NBP11 heavy chain framework 3 sequence is derived from the IGHV4-34 germline.

## References

1. WHO (2020) WHO Coronavirus Disease (COVID-19) Dashboard. https://covid19.who.int/.

2. Hoffmann M, Kleine-Weber H, Schroeder S, Krüger N, Herrler T, Erichsen S, Schiergens TS, Herrler G, Wu NH, Nitsche A, et al. (2020) SARS-CoV-2 Cell Entry Depends on ACE2 and TMPRSS2 and Is Blocked by a Clinically Proven Protease Inhibitor. Cell 181:271–280.e8.

3. Andersen KG, Rambaut A, Lipkin WI, Holmes EC, Garry RF (2020) The proximal origin of SARS-CoV-2. Nature Medicine 26.

4. Anderson DE, Tan CW, Chia WN, Young BE, Linster M, Low JGH, Tan YJ, Chen MIC, Smith GJD, Leo YS, et al. (2020) Lack of cross-neutralization by SARS patient sera towards SARS-CoV-2. Emerging Microbes and Infections 9:900–902.

5. Lv H, Wu NC, Tsang OTY, Yuan M, Perera RAPM, Leung WS, So RTY, Chan JMC, Yip GK, Chik TSH, et al. (2020) Cross-reactive Antibody Response between SARS-CoV-2 and SARS-CoV Infections. Cell Reports.

6. Ju B, Zhang Q, Ge J, Wang R, Sun J, Ge X, Yu J, Shan S, Zhou B, Song S, et al. (2020) Human neutralizing antibodies elicited by SARS-CoV-2 infection. Nature [Internet] 584:115–119. Available from: http://www.nature.com/articles/s41586-020-2380-z

7. Robbiani DF, Gaebler C, Muecksch F, Lorenzi JCC, Wang Z, Cho A, Agudelo M, Barnes CO, Gazumyan A, Finkin S, et al. (2020) Convergent antibody responses to SARS-CoV-2 in convalescent individuals. Available from: https://doi.org/10.1038/s41586-020-2456-9

8. Nielsen SCA, Yang F, Jackson KJL, Pinsky BA, Blish CA, Boyd SD, Hoh RA, Rö Ltgen K, Jean GH, Stevens BA, et al. (2020) Human B Cell Clonal Expansion and Convergent Antibody Responses to SARS-CoV-2. Available from: https://doi.org/10.1016/j.chom.2020.09.002

9. Ni L, Ye F, Cheng ML, Feng Y, Deng YQ, Zhao H, Wei P, Ge J, Gou M, Li X, et al. (2020) Detection of SARS-CoV-2-Specific Humoral and Cellular Immunity in COVID-19 Convalescent Individuals. Immunity.

10. Barnes CO, West AP, Huey-Tubman KE, Hoffmann MAG, Sharaf NG, Hoffman PR, Koranda N, Gristick HB, Gaebler C, Muecksch F, et al. (2020) Structures of Human Antibodies Bound to SARS-CoV-2 Spike Reveal Common Epitopes and Recurrent Features of Antibodies. Cell [Internet] 182:828–842.e16. Available from: https://linkinghub.elsevier.com/retrieve/pii/S0092867420307571

11. Seydoux E, Homad LJ, MacCamy AJ, Parks KR, Hurlburt NK, Jennewein MF, Akins NR, Stuart AB, Wan YH, Feng J, et al. (2020) Analysis of a SARS-CoV-2-Infected Individual Reveals Development of Potent Neutralizing Antibodies with Limited Somatic Mutation. Immunity.

12. Röltgen K, Powell AE, Wirz OF, Stevens BA, Hogan CA, Najeeb J, Hunter M, Wang H, Sahoo MK, Huang C, et al. (2020) Defining the features and duration of antibody responses to SARS-CoV-2 infection associated with disease severity and outcome. Science Immunology 5.

13. Cao Y, Su B, Guo X, Sun W, Deng Y, Bao L, Zhu Q, Zhang X, Zheng Y, Geng C, et al. (2020) Potent Neutralizing Antibodies against SARS-CoV-2 Identified by High-Throughput Single-Cell Sequencing of Convalescent Patients’ B Cells. Cell [Internet] 182:73–84.e16. Available from: https://linkinghub.elsevier.com/retrieve/pii/S0092867420306206

14. Gao Q, Bao L, Mao H, Wang L, Xu K, Yang M, Li Y, Zhu L, Wang N, Lv Z, et al. (2020) Development of an inactivated vaccine candidate for SARS-CoV-2. Science:eabc1932.

15. Ravichandran S, Coyle EM, Klenow L, Tang J, Grubbs G, Liu S, Wang T, Golding H, Khurana S (2020) Antibody signature induced by SARS-CoV-2 spike protein immunogens in rabbits. Science Translational Medicine:eabc3539.

16. Lambe T, Spencer A, Belij-Rammerstorfer S, Purushotham JN, Port JR, Avanzato V, Bushmaker T, Ulaszewska M, Feldmann F, Allen ER, et al. ChAdOx1 nCoV-19 vaccination prevents SARS-CoV-2 pneumonia in rhesus macaques Neeltje van Doremalen. Available from: https://doi.org/10.1101/2020.05.13.093195

17. Du L, Zhao G, Li L, He Y, Zhou Y, Zheng BJ, Jiang S (2009) Antigenicity and immunogenicity of SARS-CoV S protein receptor-binding domain stably expressed in CHO cells. Biochemical and Biophysical Research Communications 384:486–490.

18. Quinlan BD, Mou H, Zhang L, Guo Y, He W, Ojha A, Parcells MS, Luo G, Li W, Zhong G, et al. The SARS-CoV-2 receptor-binding domain elicits a potent neutralizing response without antibody-dependent enhancement. Available from: https://doi.org/10.1101/2020.04.10.036418

19. Rogers TF, Zhao F, Huang D, Beutler N, Burns A, He W, Limbo O, Smith C, Song G, Woehl J, et al. (2020) Isolation of potent SARS-CoV-2 neutralizing antibodies and protection from disease in a small animal model. Science [Internet]:eabc7520. Available from: https://www.sciencemag.org/lookup/doi/10.1126/science.abc7520

20. Powell AE, Zhang K, Sanyal M, Tang S, Weidenbacher PA, Li S, Pham TD, Pak JE, Chiu W, Kim PS (2021) A Single Immunization with Spike-Functionalized Ferritin Vaccines Elicits Neutralizing Antibody Responses against SARS-CoV-2 in Mice. ACS Central Science.

21. Ju B, Zhang Q, Ge J, Wang R, Sun J, Ge X, Yu J, Shan S, Zhou B, Song S, et al. (2020) Human neutralizing antibodies elicited by SARS-CoV-2 infection. Nature.

22. Premkumar AL, Segovia-Chumbez B, Jadi R, Martinez DR, Raut R, Markmann A, Cornaby C, Bartelt L, Weiss S, Park Y, et al. The RBD Of The Spike Protein Of SARS-Group Coronaviruses Is A Highly Specific Target Of SARS-CoV-2 Antibodies But Not Other Pathogenic Human and Animal Coronavirus Antibodies. Available from: https://doi.org/10.1101/2020.05.06.20093377

23. Shang J, Wan Y, Luo C, Ye G, Geng Q, Auerbach A, Li F Cell entry mechanisms of SARS-CoV-2. Available from: www.pnas.org/cgi/doi/10.1073/pnas.2003138117

24. Wang Q, Zhang Y, Wu L, Niu S, Song C, Zhang Z, Lu G, Qiao C, Hu Y, Yuen KY, et al. (2020) Structural and Functional Basis of SARS-CoV-2 Entry by Using Human ACE2. Cell 181:894–904.e9.

25. Mehta N, Maddineni S, Kelly RL, Lee RB, Hunter SA, Silberstein JL, Parra Sperberg RA, Miller CL, Rabe A, Labanieh L, et al. (2020) An engineered antibody binds a distinct epitope and is a potent inhibitor of murine and human VISTA. Scientific Reports [Internet] 10:15171. Available from: http://www.nature.com/articles/s41598-020-71519-4

26. Li F (2016) Structure, Function, and Evolution of Coronavirus Spike Proteins. Annual Review of Virology 3.

27. Yuan M, Wu NC, Zhu X, Lee C-CD, So RTY, Lv H, Mok CKP, Wilson IA (2020) A highly conserved cryptic epitope in the receptor binding domains of SARS-CoV-2 and SARS-CoV. Science [Internet] 368:630–633. Available from: http://www.sciencemag.org/lookup/doi/10.1126/science.abb7269

28. Brouwer PJ, Caniels TG, van der Straten K, Snitselaar JL, Aldon Y, Bangaru S, Torres JL, Okba NM, Claireaux M, Kerster G, et al. (2020) Potent neutralizing antibodies from COVID-19 patients define multiple targets of vulnerability. Science 369:643–650.

29. Barnes CO, Jette CA, Abernathy ME, Dam K-MA, Esswein SR, Gristick HB, Malyutin AG, Sharaf NG, Huey-Tubman KE, Lee YE, et al. (2020) Structural classification of neutralizing antibodies against the SARS-CoV-2 spike receptor-binding domain suggests vaccine and therapeutic strategies. bioRxiv [Internet]:2020.08.30.273920. Available from: http://biorxiv.org/content/early/2020/08/30/2020.08.30.273920.abstract

30. Crawford KH, Eguia R, Dingens AS, Loes AN, Malone KD, Wolf CR, Chu HY, Alejandra Tortorici M, Veesler D, Murphy M, et al. (2020) Protocol and reagents for pseudotyping lentiviral particles with SARS-CoV-2 Spike protein for neutralization assays. Viruses [Internet] 12:513–undefined. Available from: https://doi.org/10.1101/2020.04.20.051219

31. Cai Y, Zhang J, Xiao T, Peng H, Sterling SM, Walsh RM, Rawson S, Rits-Volloch S, Chen B (2020) Distinct conformational states of SARS-CoV-2 spike protein. Science.

32. Zhou D, Duyvesteyn HME, Chen C-P, Huang C-G, Chen T-H, Shih S-R, Lin Y-C, Cheng C-Y, Cheng S-H, Huang Y-C, et al. (2020) Structural basis for the neutralization of SARS-CoV-2 by an antibody from a convalescent patient. Nature Structural & Molecular Biology [Internet]. Available from: http://www.nature.com/articles/s41594-020-0480-y

33. Shi R, Shan C, Duan X, Chen Z, Liu P, Song J, Song T, Bi X, Han C, Wu L, et al. (2020) A human neutralizing antibody targets the receptor-binding site of SARS-CoV-2. Nature [Internet] 584:120–124. Available from: http://www.nature.com/articles/s41586-020-2381-y

34. Murin CD, Wilson IA, Ward AB (2019) Antibody responses to viral infections: a structural perspective across three different enveloped viruses. Nature Microbiology 4.

35. Zost SJ, Gilchuk P, Chen RE, Brett Case J, Trivette A, Nargi RS, Sutton RE, Chen EC, Binshtein E, Shrihari S, et al. Rapid isolation and profiling of a diverse panel of human monoclonal antibodies targeting 3 the SARS-CoV-2 spike protein 4 5. Available from: https://doi.org/10.1101/2020.05.12.091462

36. Avnir Y, Watson CT, Glanville J, Peterson EC, Tallarico AS, Bennett AS, Qin K, Fu Y, Huang C-Y, Beigel JH, et al. (2016) IGHV1-69 polymorphism modulates anti-influenza antibody repertoires, correlates with IGHV utilization shifts and varies by ethnicity OPEN. Available from: www.nature.com/scientificreports/

37. Amanat F, Stadlbauer D, Strohmeier S, O Nguyen TH, Chromikova V, McMahon M, Jiang K, Asthagiri Arunkumar G, Jurczyszak D, Polanco J, et al. (2020) A serological assay to detect SARS-CoV-2 seroconversion in humans. Available from: https://doi.org/10.1038/s41591-020-0913-5

38. Li Y, Sousa R (2012) Expression and purification of E. coli BirA biotin ligase for in vitro biotinylation. Protein Expression and Purification 82.

